# Genome-wide high-density CRISPR interference screens reveal condition-specific metabolic vulnerabilities in *Pseudomonas aeruginosa* PAO1

**DOI:** 10.1101/2025.08.12.669819

**Authors:** Andreas Kaczmarczyk, Alexander Klotz, Pablo Manfredi, Urs Jenal

## Abstract

*Pseudomonas aeruginosa* is a metabolically versatile opportunistic human pathogen. It causes acute and chronic infections and is notorious for its multidrug resistance and tolerance. To systematically uncover genetic vulnerabilities that could be exploited as therapeutic targets, we present a portable high-density CRISPR interference (CRISPRi) library comprising >80’000 single-guide RNAs (sgRNAs) targeting virtually all annotated coding sequences and intergenic regions of *P. aeruginosa* PAO1. This library was used to assess the genome-wide fitness landscapes under different growth conditions, uncovering gain- and loss-of-function phenotypes for more than a thousand genes upon depletion. Many of the phenotypes are likely caused by hypomorphic (partial loss-of-function) alleles that would not be easily accessible by traditional transposon sequencing (Tn-Seq). Focusing on central carbon metabolism, we reveal two glyceraldehyde-3-phosphate dehydrogenases as central, non-redundant nodes in glycolytic and gluconeogenic growth conditions that might be promising targets to redirect carbon flux away from metabolically persistent states associated with chronic infections. More generally, our comprehensive sgRNA libraries are a valuable resource to access genome-wide quantitative phenotypes through CRISPRi beyond the binary phenotypes offered by Tn-Seq.

## INTRODUCTION

*Pseudomonas aeruginosa* is an opportunistic bacterial pathogen responsible for severe infections in immunocompromised patients, notably those suffering from cystic fibrosis (CF), chronic obstructive pulmonary disease (COPD), burn wounds, and hospital-acquired infections (Folkesson *et al*, 2012; Martinez-García *et al*, 2022; Stover *et al*, 2000). In CF patients, *P. aeruginosa* colonization often leads to persistent, chronic infections that significantly contribute to morbidity and mortality due to *P. aeruginosa*’s intrinsic antibiotic resistance or tolerance and adaptability to diverse environmental stresses (Santi *et al*, 2021; Mulcahy *et al*, 2010; Levin-Reisman *et al*, 2017). A hallmark of *P. aeruginosa* biology is its exceptional metabolic versatility (Lohia & Riquelme, 2025). This bacterium can utilize a wide array of carbon and nitrogen sources, enabling it to thrive in diverse ecological niches, ranging from soil and water environments to complex host tissues (Diggle & Whiteley, 2020, 2021). Metabolic adaptability is closely tied to its large genome and extensive regulatory networks (Stover *et al*, 2000; Payne *et al*, 2021; Sánchez-Jiménez *et al*, 2023; Park & Sauer, 2022; Laventie & Jenal, 2020; Manner *et al*, 2023), facilitating survival under nutrient limitation, oxidative stress, and antibiotic exposure.

*P. aeruginosa* exhibits significant genetic diversity characterized by a large pangenome, which encompasses numerous accessory genes alongside a conserved core genome. This extensive genetic repertoire allows for rapid evolution and adaptation through horizontal gene transfer, mutation, and recombination, further complicating clinical management and treatment strategies (Klockgether *et al*, 2010; Freschi *et al*, 2019; Yi & Dalpke, 2022). Understanding the genetic basis of this flexibility and adaptability is crucial for developing targeted therapeutic interventions. Traditional methods for functional genomic studies in *P. aeruginosa*, such as transposon sequencing (Tn-seq) (Lee *et al*, 2015; Poulsen *et al*, 2019; Karash & Yahr, 2022; Janet-Maitre *et al*, 2023; Schinner *et al*, 2020; Skurnik *et al*, 2013; Liberati *et al*, 2006) face important limitations, including difficulties in recovering hypomorphic (partial loss-of-function) phenotypes, and limited resolution for regulatory and intergenic regions. CRISPR interference (CRISPRi), a technique enabling precise, tunable and reversible gene expression knockdown without permanent genome alterations, represents a significant methodological advancement. CRISPRi employs a catalytically inactive (“dead”) Cas9 variant (dCas9) programmed with a single-guide RNA (sgRNA) to bind specific DNA sequences with appropriate protospacer-adjacent motifs (PAMs), acting as a physical roadblock to RNA polymerase (RNAP)-mediated transcription initiation or elongation (Qi *et al*, 2013; Jinek *et al*, 2012). When targeted to the non-template (NT) strand within a coding sequence, dCas9 impedes RNAP progression, whereas binding to both the NT or the template (T) strand in promoter regions can block transcription initiation by masking sigma factor-binding sites (Qi *et al*, 2013; Hall *et al*, 2022). Generally, targeting of dCas9 to the 51-end of the coding sequence seems to be more efficient, probably due to fewer RNAP complexes trailing and colliding with the dCas9-sgRNA roadblock (Qi *et al*, 2013; Gilbert *et al*, 2014; Ran *et al*, 2015). Thus, targeting dCas9 to different positions along the coding sequence provides a wide range of repression levels, thereby allowing graded knockdown of gene expression. In addition, CRISPRi effects are reversible and adjustable by inducer concentration when dCas9 is placed under control of an inducible promoter (Qi *et al*, 2013). CRISPRi thus permits functional characterization of both essential genes and non-essential genes and regulatory elements in a highly quantitative manner, overcoming many of the limitations inherent to the binary essentialities defined by Tn-Seq.

In this study, we leverage a dual-plasmid system with chromosomally integrated dcas9 and episomal sgRNA expression (Tan *et al*, 2018). We present the development and validation of portable high-density, genome-wide CRISPRi sgRNA libraries tailored for high-throughput functional genomic studies in *P. aeruginosa* PAO1. Our studies showcase the potential of large sgRNA libraries by systematically exploring essential and condition-specific gene requirements under multiple nutritional conditions. Our findings highlight critical metabolic pathways, regulatory networks, and novel genetic elements involved in *P. aeruginosa* growth and adaptation. This high-resolution, genome-wide functional map deepens our understanding of *P. aeruginosa* biology and represents a valuable community resource that will help to identify potential targets for novel antimicrobial strategies.

## RESULTS AND DISCUSSION

### Modified expression plasmid for efficient construction of high-fidelity sgRNA libraries

In order to construct a high-density sgRNA library for *P. aeruginosa* PAO1, we chose a previously described dual-plasmid system (Tan *et al*, 2018). Here, *dcas9* from *Streptococcus pasteurianus* is chromosomally integrated at the *attTn7* locus with expression under control of the IPTG-inducible *lacIq/Plac* system derived from E. coli (Choi *et al*, 2005; Choi & Schweizer, 2006). The sgRNA is expressed episomally from a pBBR-based plasmid, pBx-Spas-sgRNA-Gm (Figure 1a). Although its transcription was originally intended to be anhydrotetracycline-inducible (via *tetR/Ptet*), expression was found to be constitutive (Tan *et al*, 2018), possibly due to readthrough transcription from the upstream *aacC1* promoter driving expression of the gentamycin resistance cassette (Figure 1a). This feature is, in fact, beneficial as it simplifies the system to require only a single inducer, IPTG. The original sgRNA expression plasmid, pBx_Spas-sgRNA-Gm, was designed for sgRNA spacer cloning by Golden Gate assembly (GGA) (Engler *et al*, 2008) via the type IIS restriction enzyme *BbsI*. Whereas this is sufficient for single spacers, cloning of large libraries of spacers requires more efficient conditions to cover the entire spacer diversity. In particular, GGA was shown to yield the largest amount of clones and high-fidelity assembly when using a single, isothermal overnight incubation step at 37 °C (Potapov *et al*, 2018). *BbsI* is not suited for this protocol since, depending on the exact product, the enzyme shows either star activity or loses activity during prolonged incubation times (www.NEB.com). We thus replaced the *BbsI* restriction sites by BsaI sites, which allows the use of BsaI-HFv2, a type IIS restriction system that remains active during prolonged incubation times and has been extensively optimized and tested for high-fidelity GGA (Sikkema *et al*, 2023; Pryor *et al*, 2020). In addition, we included a dropout cassette encoding a counterselection marker, ccdB, in between the two BsaI sites. After GGA and transformation any cell carrying the empty sgRNA expression plasmid will be eliminated due to CcdB-mediated toxicity (Bernard & Couturier, 1992; Bernard *et al*, 1994). Because similar counterselection strategies are well-established as part of Gateway cloning (Hartley *et al*, 2000), we reasoned that ccdB would greatly reduce the rate of false-positive clones in GGA, which could potentially skew pooled sgRNA library CRISPRi experiments. The final optimized plasmids were called pBx-Gm and pBx-Amp and a schematic of the respective modifications is shown in Figure 1a.

**Figure 1.**
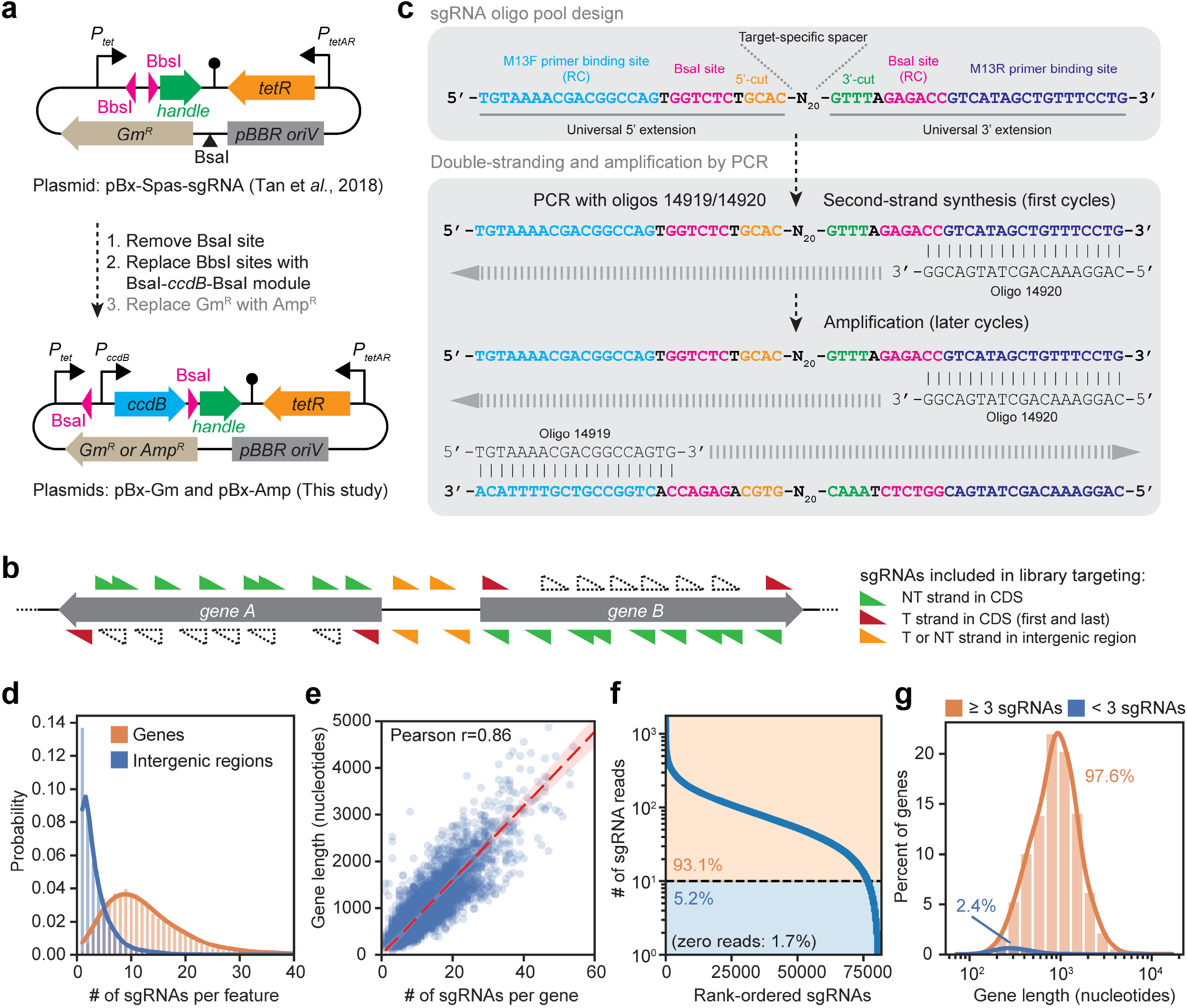
sgRNA library design and characterization. **a, Schematic of construction of pBx-Gm and pBx-Amp, two plasmids for high-efficiency, high-fidelity sgRNA cloning. b**, Schematic of the positions of PAMs for which sgRNAs were included in the library in the NT (green triangles) or T (red triangles) strand of coding sequences, or on both strands in intergenic regions (orange triangles). PAMs on the T strand for which no sgRNAs were included are shown by dotted triangles. CDS, coding sequence. **c**, Schematic of the sgRNA oligo pool design and the PCR-based second-stranding and amplification method using oligos 14919 and 14920 priming in universal flanking M13 and M13R binding regions. RC, reverse complement. **d**, Distribution of number of sgRNAs per gene or intergenic region. **e**, Correlation of number of sgRNAs per gene and gene length. **f**, Distribution of sequencing reads for all designed sgRNAs in the final pBx-Gm-based library. **g**, Distribution of gene length of genes that are covered by at least 3 sgRNAs, each with at least 10 sequencing reads (orange), and those that do not meet this criterion (blue) shown for the pBx-Gm-based library. Note that characteristics of the pBx-Amp-based library are very similar to the ones shown here for the pBx-Gm-based library.

### Design, construction and characterization of CRISPRi sgRNA libraries

Cas9 of S. pasteurianus targets protospacer adjacent motifs (PAMs) with sequences NNGTGA and NNGCGA, collectively NNGYGA (Ran *et al*, 2015; Tan *et al*, 2018). We computationally scanned the *P. aeruginosa* PAO1 genome of our laboratory strain UJP505 (see Methods) for NNGYGA PAMs and extracted adjacent 20-nt sequences corresponding to sgRNA spacers (Figure 1b). Of those, all spacers that targeted coding sequences (CDSs) in the non-template (NT) strand were included, since it was shown that dCas9-sgRNA complexes interfere with RNAP elongation more efficiently when directed against the NT strand (Qi *et al*, 2013; Hall *et al*, 2022). We also included spacer sequences targeting PAMs in both strands of intergenic regions. The rationale was that these regions might harbor promoters, and the dCas9-sgRNA complex can target promoters efficiently on both strands when directed towards the -35 or -10 core promoter elements by interfering with initial RNAP binding (transcription initiation) rather than elongation (Qi *et al*, 2013). Moreover, intergenic regions may contain unannotated coding sequences, non-coding RNAs, or other unknown features that could be amenable to targeting by CRISPRi. If possible, we also included two spacers targeting the first and the last PAM in the T strand of genes, which we rationalized would have a much milder effect on gene silencing and thus mimic weak hypomorphic alleles.

After appending sequences for amplification and cloning (Figure 1c) and additional filtering steps (see Methods), the final sgRNA oligo pool design (Supplementary Data 1) comprised 81’832 unique sgRNAs targeting the *P. aeruginosa* PAO1 genome with an average (median) of 11 sgRNAs per gene (mean +/-SD of 12.6 +/-8.1) and two sgRNAs per intergenic region (mean +/-SD of 3.2 +/-2.9) (Figure 1d). As expected, the number of sgRNAs per gene correlated well with gene length (Figure 1e). The purchased pool of oligos was double-stranded, amplified, cloned by GGA, and the resulting plasmid libraries were deep-sequenced. More than 98.3% and 98.7% of designed sgRNAs were detected in pBx-Gm- and pBx-Amp-based libraries, respectively. Moreover, over 93% of all sgRNAs were represented in the pooled libraries by at least ten sequencing reads (Figure 1f; Tables S1, S2). Almost 98% of all *P. aeruginosa* genes were covered by at least three sgRNAs with at least ten sequencing reads each (Figure 1g). As expected, genes that failed to match these criteria were strongly skewed towards smaller size, indicating that less PAMs were available. From this, we conclude that the sgRNA libraries are of good quality and likely allow robust quantification of enrichment and depletion scores on a genome-wide scale.

### High-density CRISPRi screening reveals known and novel fitness determinant

In order to validate the sgRNA libraries, we decided to set up screens for high-fitness (“essential”) genes under different growth conditions, namely rich medium represented by Lysogeny Broth (LB)-Miller, and MOPS minimal media with glucose or succinate as the sole carbon source (Figure 2). These conditions were chosen because *P. aeruginosa* prefers growth on amino acids or succinate and other TCA cycle intermediates as carbon source over growth on glucose, even when glucose is much more abundant (Folkesson *et al*, 2012; La Rosa *et al*, 2019; Palmer *et al*, 2007; Rojo, 2010; Lohia & Riquelme, 2025). On the other hand, growth in the mucus layer of human airways in some aspects resembles growth on glucose (Meirelles *et al*, 2024). This suggests that *P. aeruginosa* can flexibly adapt its metabolism during infection and that understanding the molecular underpinnings might unveil druggable targets for clinical intervention. In addition, *Pseudomonas* has an unusual central carbon metabolism, employing parts of the Embden-Meyerhoff-Parnas (EMP), the Entner-Douderoff (ED) and the Pentose Phosphate (PP) pathways, called the EDEMP cycle (Nikel *et al*, 2015; Dolan *et al*, 2020). Operation of this pathway was proposed to provide *Pseudomonas* with more reductive power in form of NADPH and might contribute to the exceptionally high resilience of this bacterium towards oxidative stress (Nikel *et al*, 2021), conditions naturally experienced during infection as part of the host immune response (Lavoie *et al*, 2011; Winterbourn *et al*, 2016). Growth on LB was included because previous studies used similar conditions and Tn-Seq to determine essential genes in *P. aeruginosa* PAO1 (Lee *et al*, 2015), thus allowing us to benchmark our dataset against these studies.

**Figure 2.**
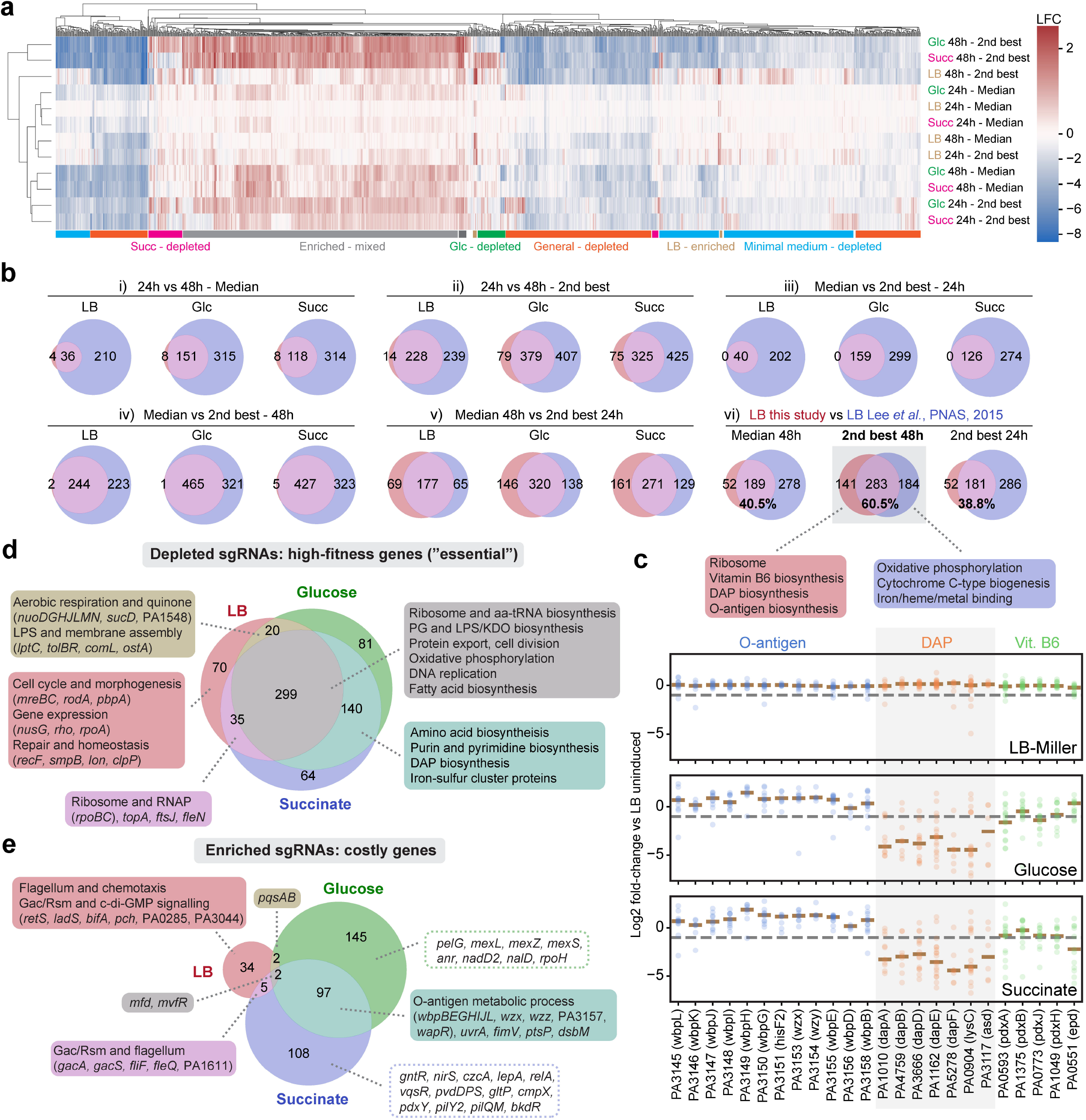
Genome-wide CRISPRi screens reveal condition-specific fitness determinants. **a**, Heatmap of log2 fold-changes (LFC) of all genes that showed significant enrichment or depletion (P-val<0.05, |LFC|>1) in at least one condition plotted using the seaborne.clustermap function and average clustering method. **b**, Venn diagrams depicting pairwise comparisons of Median and Second-best quantification methods after 24h or 48h of growth (panels i-v) and indicated comparisons to the gene essentiality data by Lee *et al*., 2015 (panel vi). Note that for panels i-v both enriched and depleted genes were included, whereas for comparisons in panel vi only depleted genes, i.e. genes whose CRISPRi-mediated repression causes fitness defects, were used. The number of genes in each sector of the Venn diagrams is indicated. **c**, Top: Functionally enriched categories in the current (red) or the Lee *et al*. dataset (blue) on LB medium as determined using the String database (Szklarczyk *et al*, 2023). Bottom: LFCs for all sgRNAs of select genes under different growth conditions. Each dot represents the LFC of a single sgRNA calculated from four independent biological replicates, and the median of all sgRNA LFCs for one gene is indicated by a brown line. The grey dashed line marks the -1 LFC threshold. For genes, PA numbers (locus tags) as well as their annotations according to a reference genome annotation file (Stover *et al*, 2000) are given at the bottom. **d**, Venn diagrams for genes for which sgRNAs were depleted in at least one of the three tested growth conditions upon induction versus the LB uninduced control are shown along with functionally enriched categories according to the String database. Single select genes are listed in parentheses if the belonged to an enriched category or without if they were not recognized as such by String. **e**, Similar to d, but for genes whose downregulation improved fitness, i.e. their sgRNAs were enriched. The dotted boxes for glucose and succinate conditions indicate that no functionally enriched categories were identified; instead, we highlight individual genes that deemed interesting to us.

*P. aeruginosa PAO1* strains expressing chromosomal *dcas9* were transformed with the pBx-Gm-based sgRNA library and cultured in LB without induction, followed by sub-culturing into LB, MOPS/glucose, or MOPS/succinate with IPTG induction; non-induced LB cultures served as controls. Samples were sequenced after 24h and 48h (with re-dilution at 24h), which roughly corresponds to 7-8 and 14-16 generations, and analyzed using the MAGeCK pipeline (Li *et al*, 2014). Two built-in quantification methods were compared: “Median” (median sgRNA log2 fold-change [LFC] of all sgRNAs) and “Second-best” (second-best sgRNA LFC). The latter method formally corresponds to the median of the three best sgRNAs assuming at least three sgRNAs per gene. “Median” is conservative, reducing false positives but increasing false negatives. This is especially the case for longer genes because sgRNAs targeting the 3’-part of genes far from the transcriptional start site are expected to be less efficient (Qi *et al*, 2013; Gilbert *et al*, 2014). Conversely, “Second-best” is more sensitive and expected to capture more genes. Clustering of depletion/enrichment profiles (Figure 2a; Tables S3, S4) revealed genes essential in all media, in minimal media only, or specific to glucose or succinate, plus genes whose knockdown conferred a fitness advantage. At 24h, “Median” identified few essential genes (e.g., 40 genes in LB), whereas “Second-best” detected two to five times more depending on the condition; by 48h both sets expanded, and virtually all hits of the 24h time point persisted (Figure 2b, panels i and ii). Essentially all genes identified by the “Median” method are included in the “Second-best” sets (Figure 2b, panels iii and iv). A substantial overlap between the data sets recorded at 24h by “Second-best” and at 48h by “Median” indicates a high true-positive rate for the “Second-best” method (Figure 2b, panel v). When grown in LB, the 48h “Second-best” set overlapped to about 60% with essential genes determined by Tn-Seq in a previous study (Lee *et al*, 2015). In contrast, only roughly 40% of the 48h “Median” or 24h “Second-best” sets overlapped with the Tn-Seq data (Figure 2b, panel vi), arguing that “Second-best” clearly outperforms “Median”. The discrepancies between our results and those by Lee *et al*. may reflect incomplete CRISPRi knockdown (i.e. penetrance) or differences in media composition and strain genotype. For instance, diaminopimelic acid (DAP) and vitamin B6 biosynthesis are essential in the dataset from Lee *et al*., but dispensable in our LB media conditions, yet they become essential in minimal media (Figure 2c-d). This suggests that i) our LB formulation contains sufficient amounts of these compounds to support growth of auxotrophs, and ii) that the sgRNA libraries are able to capture the essentiality of these genes under appropriate conditions. From this, we concluded that the 48h “Second-best” metric offers the most inclusive and sensitive definition of essential genes and it was thus adopted for subsequent enrichment analyses. Our results also suggest that – with large sgRNA libraries and without a *priori* knowledge of the effectiveness of individual sgRNAs – an sgRNA-centric quantification approach (e.g., “Second-best”) performs substantially better than a gene-centric approach (e.g., “Median”).

### General fitness effects

The genome-wide CRISPRi screen in *P. aeruginosa* PAO1 identified a broad set of genes, the knockdown of which severely impaired growth under all conditions tested (299 genes, Figure 2d), highlighting core cellular processes essential for fitness. As expected, many of the highest-confidence hits were genes involved in fundamental cellular processes and metabolism. These include cell division genes (*fts* genes), peptidoglycan biosynthesis genes (*mur* loci and *ddlB*), DNA replication (*dnaA* and *dnaN*), and protein secretion machinery (*sec* genes, *ffh*). Multiple cell envelope biogenesis genes were also essential, such as those for lipid A and LPS biosynthesis (*lpx* enzymes), LPS transport (*msbA*), and lipoprotein trafficking (*lol* pathway). Additionally, genes for cytochrome C-type maturation (*ccm* operon), ubiquinone synthesis (*ubi* genes), heme biosynthesis (*hem* genes) and fatty acid synthesis (*fab* genes) were required for robust growth. Beyond these conserved essentials (Grazziotin *et al*, 2015; Peters *et al*, 2016), our screen exposed several *P. aeruginosa*-specific vulnerabilities. Notably, depletion of the *gatA, gatB*, and *gatC* genes, encoding subunits of the GatABC tRNA amidotransferase, caused pronounced fitness defects. *P. aeruginosa* lacks a canonical asparaginyl-tRNA synthetase, instead relying on GatABC to synthesize Asn-tRNA^Asn^ from Asp-tRNA^Asn^ (Akochy *et al*, 2004). Thus, knockdown of *gatABC* likely disrupts protein synthesis. The iron uptake regulator Fur also emerged as critical for growth. While not essential in many bacteria, *fur* in *P. aeruginosa* is difficult to delete and appears conditionally essential (Pasqua *et al*, 2017), possibly because its loss leads to the dysregulation of iron homeostasis. Another striking hit was *prtR*, which encodes a ROS-inducible repressor of pyocin bacteriocin genes (Sun *et al*, 2014). Possibly *prtR* knockdown triggers uncontrolled expression of pyocin toxins without proper immunity, resulting in self-inflicted cellular damage (Penterman *et al*, 2014). Finally, the F_1_F_0_-ATP synthase subunits (*atp* operon) seemed essential under the conditions tested. This is consistent with the respiratory preference of *P. aeruginosa* for energy production, even though it has some fermentative capabilities (Glasser *et al*, 2014; Eschbach *et al*, 2004; Tokunou *et al*, 2022). The results are consistent with the essential nature of basic cellular processes in *P. aeruginosa*, largely mirroring essential gene sets observed in other bacteria, while also pinpointing vulnerabilities specific to *P. aeruginosa*. Overall, these data demonstrate the functionality of our CRISPRi library for genome-wide fitness profiling.

Interestingly, our screens also revealed a subset of genes for which CRISPRi-mediated repression conferred a growth advantage, suggesting the targeted genes impose a burden under these conditions (Figure 2e). For example, targeting the quorum signaling genes *pqsA, pqsB*, and the PQS regulon activator (*pqsR*, also called *mvfR*) improved the relative fitness of *P. aeruginosa*. This suggests that active production of PQS signal molecules or downstream secondary metabolites might be metabolically costly or detrimental *in vitro* (Diggle *et al*, 2007). Indeed, quorum-sensing “cheater” mutants that do not produce public goods often have a growth advantage, supporting the idea that repressing pqs genes frees resources for growth (Wilder *et al*, 2011). Similarly, repression of *mfd*, which encodes the transcription-repair coupling factor, also seems to enhance fitness. Mfd detects stalled RNAP at DNA lesions, removes the stalled RNAP from DNA, and recruits the nucleotide excision repair (NER) machinery via UvrA interaction to repair the DNA lesion (Selby, 2017). Mfd can also remove RNAP stalled at protein roadblocks, such as LacI or CodY (Belitsky & Sonenshein, 2011; Hao *et al*, 2014). CRISPRi relies on the dCas9-sgRNA complex acting as a strong roadblock for RNAP for its sgRNA-defined target (Qi *et al*, 2013). In order to identify the target, hundreds of those complexes scan the genome to probe thousands of PAMs for sgRNA complementarity at any given time, representing thousands of transient roadblocks that could potentially obstruct RNAP processivity. It is thus possible that reducing Mfd activity in a CRISPRi context alleviates energy-intensive DNA repair processes or limits potentially deleterious conflicts between transcription and replication on a genome-wide scale, thereby improving growth rates.

### Fitness effects in minimal media

In defined minimal medium, we observed that many biosynthetic pathways were critical for growth, revealing numerous conditionally essential genes (140 genes, Figure 2d). As expected, auxotrophies were unmasked for amino acid and nucleotide biosynthesis (*pur* and *pyr* genes), as well as for the diaminopimelate (DAP) pathway, which produces meso-diaminopimelate for peptidoglycan cross-linking and lysine biosynthesis. Genes involved in cofactor production also showed medium-specific essentiality, including several iron-sulfur (Fe-S) cluster assembly factors and vitamin biosynthesis enzymes (*e*.*g*., biotin, riboflavin, and thiamine). The screen also highlighted genes of the central carbon metabolism that are essential only on specific minimal substrates, which will be discussed separately below. We also observed distinct sets of genes shared between growth on glucose and LB (20 genes, Figure 2d), or between succinate and LB (35 genes, Figure 2d). While the former group includes genes associated with LPS biogenesis and aerobic respiration (*e*.*g*., the nuo operon), the latter shows no major enriched functional categories except that some transcription- and translation-associated genes are overrepresented.

Similar to growth on rich medium, the minimal medium screen uncovered a set of genes whose repression enhanced fitness under nutrient limitation. This included genes associated with extracellular structures and virulence factors. For example, repression of the *wbp* gene cluster involved in O-antigen LPS synthesis (Rocchetta *et al*, 1999) improved fitness in minimal media (Figure 2e), an effect that is also seen using the more conservative “Median” comparison method (Figure 2c). This observation is somewhat surprising given that O-antigen biosynthesis was described as essential for growth in LB in a previous Tn-Seq screen (Lee *et al*, 2015). The apparent discrepancy might be explained by incomplete penetrance of CRISPRi – equivalent to a hypomorphic mutation – and a specific benefit in minimal medium due to competition between the O-antigen pathway and anabolic processes for shared sugar precursors in these more nutrient poor media (Bertani & Ruiz, 2018). Similarly, downregulating pslE, which forms part of the PSL exopolysaccharide (EPS) biofilm matrix biosynthesis operon (Byrd *et al*, 2009) conferred a growth advantage, consistent with the idea that EPS matrix production diverts resources away from growth. We also observed that guides targeting components of the Type III secretion system (T3SS) (Horna & Ruiz, 2021) and Type VI secretion system (T6SS) (Chen *et al*, 2015) were enriched in minimal medium, indicating that suppressing these virulence secretion machineries may be beneficial when nutrients are scarce. Knockdown of phenazine biosynthesis genes, which direct the production of redox-active secondary metabolites (Pierson & Pierson, 2010), and of type IV pilus assembly genes (Webster *et al*, 2022), likewise increased fitness in minimal conditions. It is plausible that these virulence structures and secondary metabolite pathways are costly in the absence of host interaction pressures or even detrimental to optimal growth in broth. Taken together, the identification of conditional essential genes and beneficial knockdowns in minimal media features how nutrient limitation reshapes the genetic landscape of essential functions in *P. aeruginosa*.

### Fitness effects in complex media

In nutrient-rich LB, we found a distinct group of 70 genes whose depletion impaired growth exclusively under these conditions (Figure 2d). Most map to functions that likely become rate-limiting under high biosynthetic flux, including several ribosomal proteins, the transcription elongation/termination factors NusG and Rho, and the RNAP α-subunit (Browning & Busby, 2004). SmpB, part of the SmpB/tmRNA rescue system (Müller *et al*, 2021), and the replication-restart protein RecF (Cox *et al*, 2023) also proved to be important under these conditions, a pattern that suggests increased translational stalling and replication stress during rapid proliferation. Cell shape maintenance emerged as another process that is specifically critical in rich media. This includes the actin homologue MreB, penicillin-binding protein PbpA and the RodA transglycosylase (Rohs & Bernhardt, 2021). Increased translational throughput during fast growth in LB also appears to raise the burden on the protein quality control system as depletion of the ATP-dependent proteases Lon and ClpP (Yang & Lan, 2016) reveals a significant fitness insult. Collectively, genes that were depleted exclusively during growth on LB indicate that even a modest reduction in transcription, translation, envelope synthesis, or proteostasis can have deleterious effects on competitive fitness during rapid growth of *P. aeruginosa*.

The screen also identified a cluster of genes, which, when depleted, led to enhanced growth, implying that their expression imposes an energetic or regulatory cost under these conditions (Figure 2e). Intriguingly, this included three major and well-defined regulatory modules. First, several genes involved in flagellar biogenesis (C-ring components FliGMN, export-gate protein FliO, ATPase FliI, and hook protein FlgK) (Ma *et al*, 2020) were strongly enriched among the group of beneficial knockdowns. Possibly, flagellar assembly consumes a substantial fraction of the cells’ energy while offering little benefit in liquid cultures. Second, components of the Gac/Rsm pathway (Lapouge *et al*, 2008; Broder *et al*, 2016; Goodman *et al*, 2004, 2009) seemed costly when fully expressed. This includes the two-component sensor kinase GacS and the response regulator GacA, together with the RetS-modulatory hybrid sensor PA1611 (Bhagirath *et al*, 2017; Kong *et al*, 2013; Chambonnier *et al*, 2016) (Figure 2e). It is possible that Gac-mediated signalling promotes the expression of factors that are redundant or detrimental during planktonic growth in LB. Third, sgRNAs targeting several c-di-GMP-specific phosphodiesterases (PDEs; *bifA, pch/dipA*, and PA0235/*pipA*) (Jenal *et al*, 2017; Römling *et al*, 2013) were enriched, together with the knockdown of the c-di-GMP-dependent global transcription regulator FleQ (Hickman & Harwood, 2008). Depletion of PDEs is expected to elevate intracellular c-di-GMP levels, and a deletion of *fleQ* was previously shown to result in a wrinkly phenotype, typically associated with strong biofilm formation similar to mutants overproducing c-di-GMP. The *fleQ* null phenotype was linked to increased production of the two major extracellular matrix (ECM) constituents Pel and Psl (Hickman & Harwood, 2008). It is thus tempting to speculate that the genes in this set share the general property of tilting regulatory networks towards a sessile, biofilm-associated state. Whether this represents a bias during our growth or sampling conditions or, indeed, has some physiological relevance remains to be shown in the future.

### Central carbon metabolism during growth in glucose or succinate

*P. aeruginosa* relies on a flexible central carbon metabolism to utilize a diverse range of substrates (Dolan *et al*, 2020; Rojo, 2010; La Rosa *et al*, 2019). Instead of classical glycolytic pathways, *Pseudomonas* uses a hybrid EDEMP cycle, combining segments of the EMP, ED and PP pathways (Figure 3a), likely to meet the organism’s high demand for NADPH (Nikel *et al*, 2015, 2021; Dolan *et al*, 2020). We find that many of the genes in central carbon metabolism show condition-dependent fitness scores (Figure 3b). within the EDEMP cycle the central enzymes transketolase (TktA), fructose-1,6-bisphosphate (FBP) aldolase (Fda), fructose-1,6-bisphosphatase (Fbp), ribulose-phosphate 3-epimerase (Rpe), and triose-phosphate isomerase (TpiA) were essential under all conditions. This is consistent with the EMP and non-oxidative PP branches providing precursors for nucleotides, cell wall and membrane biogenesis due to the absence of 6-phosphogluconate dehydrogenase (Gnd) in *P. aeruginosa*, a key enzyme in the oxidative PP pathway in other bacteria. We further find a requirement of GltA (citrate synthase), SucAB (2-oxoglutarate dehydrogenase subunits) and Sdh (succinate dehydrogenase complex) in the tricarboxylic acid (TCA) cycle in all conditions, and of pyruvate dehydrogenase (*aceE, aceF* and *lpdG*), enolase (*eno*) and phosphoglycerate mutase (*pgm*) in the lower glycolytic pathway connecting the EDEMP cycle with the TCA cycle. Production of acetyl-CoA by pyruvate dehydrogenase not only represents the key irreversible step linking the ED branch and the TCA cycle, but also provides building blocks for fatty acid, lipid and isoprenoid biosynthesis. Eno and Pgm catalyze the two-step conversion of 3-phosphoglycerate (3PG) to phosphoenolpyruvate (PEP), a unique, central and reversible reaction that is crucial for both glycolytic and gluconeogenic pathways.

**Figure 3.**
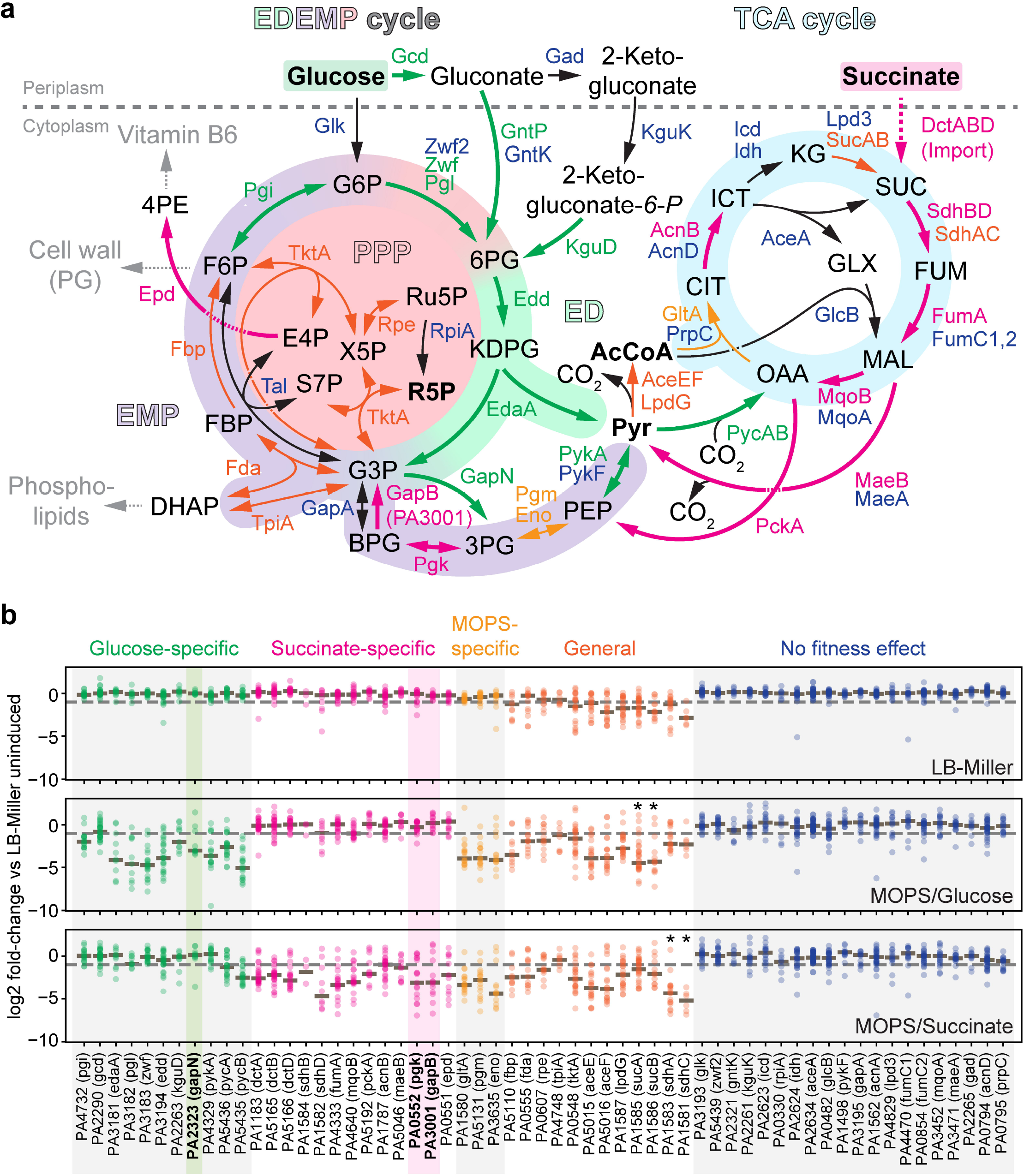
Carbon source-specific fitness determinants in central carbon metabolism. **a**, Schematic of the central carbon metabolism of *P. aeruginosa* PAO1. Metabolites are shown in black, enzymes and the reactions they potentially catalyze (arrows) are color-coded according to their CRISPRi scores: blue, no significant change under any conditions; dark orange, essential under all growth conditions; light orange, essential under minimal medium conditions in general; green, essential in glucose conditions only; magenta, essential in succinate conditions only. Parts of the EDEMP cycle that are derived from the EMP, ED or PP pathways are highlighted by different colors, as is the TCA cycle (light blue). Abbreviations for metabolites are defined in the main text or are as follows: DHAP, dihydroxyacetone phosphate; KDPG, 2-keto-3-deoxy-6-phosphogluconate; E4P, erythrose-4-phosphate; S7P, sedoheptulose-7-phosphate; X5P, xylulose-5-phosphate; CIT, citrate; SUC, succinate; FUM, fumarate; MAL, malate. **b**, LFCs for all sgRNAs of central carbon metabolism genes/proteins mapped in panel a are shown for the three different growth conditions: LB-Miller, MOPS/glucose and MOPS/succinate. Each dot represents the log2 fold-change (LFC) of a single sgRNA calculated from four independent biological replicates, and the median of all sgRNA LFCs for one gene is indicated by a brown line. The grey dashed line marks the -1 LFC threshold. For genes, PA numbers (locus tags) as well as their annotations according to a reference genome annotation file (Stover *et al*, 2000) are given at the bottom. Asterisks indicate genes that were generally essential as defined by our cutoff, but showed clear quantitative differences between glucose- and succinate-grown cells. GapN (PA2323) and GapB (PA3001) together with Pgk (PA0552), key enzymes in lower glycolysis which are differentially required during growth on glucose and succinate and discussed in the main text, are highlighted in bold and with green and magenta backgrounds, respectively.

In glucose-grown cells, we find that depletion of glucose dehydrogenase (*gcd*), gluconate permease (*gntP*), and 2-ketogluconate-6-phosphate dehydrogenase (*kguT*) show fitness defects, in agreement with glucose assimilation in *P. aeruginosa* depending on periplasmic oxidation of glucose rather than the transport-coupled glucose phosphorylation pathway known from, e.g., *E. coli* (Raneri *et al*, 2018; Nikel *et al*, 2015; Carreón-Rodríguez *et al*, 2023). While glucose flux was shown to primarily proceed via gluconate in *P. putida* (Nikel *et al*, 2015), our data suggest that in *P. aeruginosa* a substantial fraction of gluconate is further oxidized to 2-ketogluconate before entering the ED pathway. Core genes in the ED branch of the EDEMP cycle were strictly required for growth on glucose, but not succinate, including *zwf* (glucose-6-phosphate dehydrogenase), *pgl* (6-phosphogluconolactonase), *edd* (6-phosphogluconate dehydratase) and *edaA* (KDPG aldolase). This is consistent with a lack of the key EMP enzyme 6-phosphofructo-1-kinase (Pfk) in *Pseudomonas* and, consequently, hexoses being primarily catabolized via the ED branch. However, we also see a fitness effect of G6P isomerase (*pgi*) depletion, suggesting that cell wall biosynthesis and non-oxidative PP reactions are fed via this anabolic EMP route during growth on glucose. In lower glycolysis, we see a striking dependency on a single NADP^+^-dependent, non-phosphorylating glyceraldehyde-3-phosphate dehydrogenase (GAPDH), GapN (PA2323), whereas depletion of the two canonical NAD-dependent GAPDHs, *gapA* (PA3195) and PA3001 (here called *gapB*) showed no effect with glucose as the sole carbon source. GapN, by homology, is predicted to convert glyceraldehyde-3-phosphate (G3P) directly to 3-phosphoglycerate (3PG), while generating NADPH and bypassing the ATP-forming phosphoglycerate kinase (Pgk) step (Habenicht, 1997). This metabolic bypass could help to maintain redox balance by providing NADPH for anabolic, rather than NADH for catabolic pathways. Finally, flux into the TCA cycle relied on *pykA* (pyruvate kinase A) and *pycAB* (pyruvate carboxylase), but not *pykF*. The need for pyruvate carboxylase likely reflects a need for anaplerotic oxaloacetate (OAA) when glycolytic flux accelerates.

Growth on succinate required the C4-dicarboxylate transporter DctA and its regulators DctB and DctD (Valentini *et al*, 2011), and nearly every reaction in the TCA cycle became essential, in agreement with the TCA cycle’s role as the major catabolic source under these conditions. The exception was oxidative carboxylation of isocitrate (ICT) to α-ketoglutarate (KG), probably due to redundancies of Idh and Icd. Other steps with multiple possible isozymes showed clear dependencies on a single gene, e.g. *fumA* for fumerase and *mqoB* for malate dehydrogenase, highlighting their non-redundant roles during growth on succinate. Phosphoenolpyruvate carboxylase (PckA) and NADP^+^-dependent malic enzyme (MaeB) were also essential, underscoring their major roles in shuffling carbon into gluconeogenesis and providing acetyl-CoA for fatty acid, lipid and isoprenoid biosynthesis. In the lower gluconeogenic pathway, two enzymes are specifically required for growth on succinate, but not glucose: phosphoglycerate kinase (*pgk*) and the NAD^+^-dependent GAPDH GapB (PA3001). These enzymes catalyze the sequential conversion of 3PG through 1,3-bisphosphoglycerate (BPG) to G3P, a sequence that is bypassed in glucose-grown cells by the NADP^+^-dependent, non-phosphorylating GAPDH GapN. Although GapA depletion produced no phenotype in any condition tested, chromosomal proximity to prototypical ED pathway genes of both *gapA* (*edd* and *glk*) and *gapN* (*gntP* and *gntK*) suggests that both GAPDHs might operate in parallel under glycolytic conditions, with *gapN* serving additional redox-related functions. In contrast, GapB is likely solely responsible for gluconeogenic growth on succinate. Notably, *gapB* is located adjacent to PA3000 (*aroP1*), encoding a predicted amino acid transporter, reinforcing the link to gluconeogenic growth, and the nqr gene cluster (PA2994-PA2999), encoding one of three NADH dehydrogenases in *P. aeruginosa* PAO1, and the only one besides complex I (Nuo) that pumps ions (Torres *et al*, 2019; Hreha *et al*, 2021). In this context, it is worth mentioning that among the strongest hits in succinate conditions are genes for a major sodium/proton antiporter (*shaABC*, also called *mrpABC*; PA1554-PA1556) (Foreman *et al*, 2021), perhaps reflecting different requirements in the proton motive force during respiration on the two carbon sources.

Overall, our results reveal distinct sets of enzymes required for glycolytic and gluconeogenic growth. In particular, the dichotomy in lower glycolysis imposed by GapN/Pgk and GapB might offer a unique opportunity to interfere with the directionality of flux in the central carbon metabolism of *P. aeruginosa* PAO1 and other bacteria (Casas-Román *et al*, 2024; Fillinger *et al*, 2000; Omumasaba *et al*, 2004; Purves *et al*, 2010), possibly allowing to redirect metabolic flux towards a non-persistent, more vulnerable metabolic state that is more responsive to antibiotic treatment and other clinical interventions.

## CONCLUDING REMARKS

In this study, we present a comprehensive CRISPRi library targeting virtually all annotated open reading frames and intergenic regions of *P. aeruginosa* strain PAO1. This resource enables systematic, tunable repression of both coding and non-coding elements across the genome, and provides a powerful tool for high-resolution functional genomics in this metabolically versatile and clinically relevant organism. For the analyses reported here, we focused exclusively on protein-coding sequences and deliberately simplified the analysis by excluding intergenic regions and ignoring polar effects. While this allowed us to benchmark the performance of the library against classical transposon insertion sequencing (Tn-Seq) datasets, future more fine-grained analyses will aim at exploiting the full potential of our high-density sgRNA library, *e*.*g*., by using polar effects to reveal operon boundaries, gene order dependencies, or non-coding regulatory elements. Most importantly, CRISPRi enables quantitative repression of genes, offering resolution far beyond the binary essentialities defined by Tn-Seq. For example, our study was able to reveal ribosomal proteins, RNA polymerase, elongation and termination factors, and DNA, RNA and protein repair mechanisms as rate-limiting factors during fast growth in rich medium. In addition, about one third of the significant hits correspond to genes whose downregulation improves rather than impairs fitness. We suspect that many of them mimic hypomorphic alleles, providing basic functionality while reducing costs associated with normal expression levels; such alleles are inherently difficult to isolate using Tn-Seq. Beyond benchmarking our set to published Tn-Seq data, we also uncovered new functional specializations of seemingly redundant paralogs in the TCA cycle and lower glycolysis/gluconeogenesis, most notably two GAPDH isoforms that promote growth on glucose (GapN) or succinate (GapB). In summary, we not only provide a genome-wide view of fitness determinants in different disease-relevant growth conditions, but also – and most importantly – establish comprehensive and portable sgRNA libraries as a community tool for deeper, systems-level studies of gene function and regulation in *P. aeruginosa*.

## METHODS

### Strains and growth conditions

*E. coli* DH5α or DB3.1 were used for routine cloning or cloning of *ccdB*-harboring plasmids, respectively. MegaX DH10B T1R Electrocomp™ Cells (Thermo Fisher Cat#C640003) were used for library preparations (see details below). *P. aeruginosa* PAO1 strain UJP505 (GenBank: CP191349.1) was used as the wild-type strain. LB-Miller (10 g/L tryptone, 5 g/L yeast extract, 10 g/L NaCl) was used as standard growth medium for E. coli and *P. aeruginosa*. LBnoSalt (10 g/L tryptone, 5 g/L yeast extract) was used for preparation of electro-competent *P. aeruginosa* cells, following preparation and electroporation protocols described previously (Klotz *et al*, 2023). When appropriate, antibiotics were used at the following concentrations for E. coli and *P. aeruginosa*, respectively: oxytetracycline 12.5 and 100 mg/L; carbenicillin 100 and 100 mg/L; gentamycin 20 and 30 mg/L. Isopropyl b-D-thiogalactoside (IPTG) was used at a final concentration of 1 mM if not stated otherwise.

### Plasmid and strain constructions

Cloning and molecular biology was performed according to standard techniques using Phusion polymerase in GC buffer for PCR, and restriction enzymes, ligase and other DNA modifying enzymes from New England Biolabs (NEB). Gibson Assembly, Golden Gate Assembly or classic restriction/ligation cloning were used as appropriate. Oligonucleotides were ordered from Sigma-Aldrich or Microsynth (Balgach, Switzerland). Oligo phosphorylation and annealing was performed as described before (Chen *et al*, 2018; Klotz *et al*, 2023). Markerless integrations at the attTn7 locus were constructed as previously described using helper plasmids pTns2 and pFLP2 (Choi *et al*, 2005; Choi & Schweizer, 2006; Hoang *et al*, 1998). Oligonucleotides, plasmids and strains used in this work are listed in Table S5.

### Construction of sgRNA expression plasmids

The gene *ccdB* encoding a counterselection marker for pBx-Spas-sgRNA-Gm was obtained from pGGASG2, a sgRNA expression plasmid for *Caulobacter crescentus*, the construction of which is described in the following. First, for construction of pGGASG, two fragments were PCR-amplified from pBXMCS-2 Pconstitutive-sgRNA(Sth3)_ctrA (Addgene plasmid # 133339) (Guzzo *et al*, 2020) using oligo pairs 14872/14875 and 14874/14877 and a third fragment carrying ccdB was PCR-amplified from pQYD (Addgene plasmid # 48104) (Kaczmarczyk *et al*, 2013) using oligos 14873/14876. The three fragments were assembled via Golden Gate cloning using 1000 units T4 DNA ligase (NEB #M0202S) and 30 units BbsI-HF (NEB #R3539S) in T4 DNA Ligase buffer (NEB #B0202) in a total volume of 50 μl and incubation at 37°C for 4h. Then, to eliminate an internal BsaI restriction site in *ccdB*, two fragments were PCR-amplified from pGGASG using oligo pairs 15070/15073 and 15071/15072 and cloned in pGGASG digested with MluI/EcoRI using Gibson assembly (Gibson *et al*, 2009), resulting in plasmid pGGASG2. The single BsaI restriction site of pBx-Spas-sgRNA-Gm (Tan *et al*, 2018) was eliminated by PCR-amplification of the plasmid with oligos 14480/14881, digestion with DpnI to remove the template and transformation into chemo-competent DH5a, generating plasmid pBx(Bsa-). To construct plasmid pBx-Gm, *ccdB* was PCR-amplified from pGGASG2 with oligos 15012/15013 and cloned into pBx(Bsa-) by Golden Gate assembly using 1000 units T4 DNA ligase (NEB #M0202S) and 30 units BbsI-HF (NEB #R3539S) in 0.5x T4 DNA Ligase buffer (NEB #B0202) and 0.5x CutSmart buffer (NEB #B6004S) in a total volume of 50 μl. Incubation was 90 min at 37°C, 15 min at 16°C (2 cycles), 15 min at 55°C and 20 min at 80°C. pBx-Amp was constructed by PCR-amplification of *ampR* using oligos 15428/15355 and the pBx-Gm backbone using 15352/15353 and Gibson assembly of the two fragments. Note that *ampR* harbors a BsaI site that might interfere with proper Golden Gate assembly, although, practically, we note that library construction with this plasmid yielded similarly high-quality sgRNA libraries as with the pBx-Gm plasmid with essentially identical sgRNA coverage and diversity (Table S1).

### Design of a genome-wide sgRNA library for *P. aeruginosa* PAO1

Using custom Perl scripts, 20-nt-long sgRNA spacer sequences targeting all PAMs (NNGYGA; Y being T or C) in the non-template (NT) strand of annotated open-reading-frames or in any of the two strands in intergenic regions were extracted from the genome of strain UJP505, our lab strain wild-type *P. aeruginosa* PAO1 (GenBank: CP191349.1); in addition, we included spacer sequences targeting the first and the last PAM in the template (T) strand of each open-reading-frame. Off-target potential was assessed by sliding a 9-nt window within each sgRNA across both DNA strands of the genome and counting occurrences of partially matched PAMs. Candidates with multiple hits were flagged and deprioritized. Non-unique sgRNAs appearing more than once in the genome were excluded. 5’-extensions encoding the M13 primer sequence, a forward-pointing BsaI restriction site and an appropriate sequence compatible with the “promoter overhang” in pBx-Gm and pBx-Amp plasmids were added. 3’-extensions were added that contained an appropriate sequence compatible with the “sgRNA handle overhang” in pBx-Gm and pBx-Amp plasmids, a reverse-complementary BsaI site (backward-pointing) and a sequence reverse complementary to the M13R primer sequence. Together, these features would allow double-stranding of the oligo, PCR-amplification of the resulting dsDNA product, and subsequent BsaI-based Golden Gate Assembly in target plasmids pBx-Gm and pBx-Amp. From this sequence list, we excluded oligos that harbored internal BsaI sites, i.e. sites either partially or fully overlapping the sgRNA spacer part of the oligo. The final oligo pool also included approximately 3000 oligos that were not targeted to *P. aeruginosa* PAMs, but were instead designed for other purposes; importantly, these additional oligos harbored 5’- and 3’-extensions not compatible with the cloning procedures described here, so are not part of our sgRNA libraries. The final oligo pool was comprised of 83’872 individual oligos, of which 81832 were directed against the PAO1 genome. The oligo pool (CustomArray Oligo Pool-92K) was ordered from GenScript.

### Construction of a genome-wide sgRNA library for *P. aeruginosa* PAO1

The oligo pool was double-stranded and amplified using oligos 14919/14920 with Phusion polymerase in GC buffer with 0, 3 or 6% DMSO and the following cycling conditions: 30 sec at 98°C; 10 sec at 98°C, 10 sec at 60°C, 15 sec at 72°C (30 cycles); 1 min at 72°C, hold at 12°C. Different DMSO concentrations were used since DMSO is well known to affect DNA secondary structures during PCR, so we anticipated that different concentrations of DMSO would average out the bias introduced by the variable spacer regions of sgRNAs. All three PCR reactions yielded bands at the expected size, and PCR products were gel-purified (MN NucleoSpin Gel & PCR Clean-up Mini kit) and pooled, with a final DNA concentration of 16 ng/μl. The sgRNA library was then cloned in pBx-Gm or pBx-Amp using Golden Gate assembly in a reaction containing 10 μl pBx_Gm (67 ng/μl) or pBx_AmpR (72 ng/μl), 10 μl sgRNA library PCR product (16 ng/μl), 4000 units of T4 DNA ligase (NEB #M0202S) and 120 units of BsaI-HFv2 (NEB #R3733S) in ligase buffer (NEB #B0202) in a total volume of 200 μl. The reactions were incubated overnight at 37°C, purified using a NucleoSpin Gel & PCR Clean-up Mini kit (MN) with elution in 2 × 15 μl ddH_2_O, and the entire purified reactions were transformed in E. coli MegaX cells (diluted three-fold in cold ddH_2_O) by electroporation using a Bio-Rad GenePulser with the following settings: 25 μF, 400⍰Ω, 1.8⍰kV. Immediately after the pulse 900 μl of SOC were added, cells were incubated for 2h at 37°C, then transferred into 500 ml of LB-Miller containing gentamycin or carbenicillin. 5 µl were plated on selective plates to estimate library size and the remaining culture was grown in a flask at 37°C with shaking at 180 rpm. The next day, cryo-stocks of libraries were prepared and plasmids were prepped using a GenElute™ Plasmid Miniprep Kit (Sigma-Aldrich). The final pBx-Gm- and pBx-Amp-based libraries consisted of approximately 3.4 × 10^6^ and 2.4 × 10^6^ individual clones, respectively, representing 57- and 40-fold coverage.

### Transformation of sgRNA libraries in *P. aeruginosa* PAO1

Four independent cultures of *P. aeruginosa* PAO1 strain UJP4737, a derivative of the wild-type strain UJ505 carrying IPTG-inducible *dcas9* (Tan *et al*, 2018) in the chromosomal *attTn7* locus, were started from single colonies in 5 ml LBnoSalt and electro-competent cells were prepared as described before (Klotz *et al*, 2023) from 4 ml of culture for each replicate. One μl of the pBx-Gm-based library was electroporated in cells as described previously, cells were recovered in 100 ml of LB, grown for 2h at 37°C with shaking (180 rpm), gentamycin was added, and growth of cultures was continued overnight in the same conditions. A small aliquot was taken at the time when gentamycin was added and plated on selective LB plates to estimate transformation efficiency. 3.2-11.6 × 10^6^ clones were obtained per transformation, covering the theoretical library size 40-to 145-fold. Cells from 5 ml of overnight cultures were collected by centrifugation and frozen at -20°C as t_0_ samples, and aliquots corresponding to 1 ml of OD_600_=1 were used to inoculate 100 ml of different media (LB, MOPS/Glc or MOPS/Succ) in 500-ml shaking flasks without and/or with 1 mM IPTG to induce *dcas9* expression. After 24h of growth at 37°C with shaking, 5 ml of cultures were pelleted and cells were frozen at -20°C as t_24h_ samples. Cultures were then re-diluted in fresh medium and grown for another 24h in the same conditions as before to further deplete condition-specific sgRNAs, and 5-ml samples were taken and frozen as t_48h_ samples as described above.

### Illumina sequencing of sgRNA libraries

Frozen *P. aeruginosa* pellets (see above) were resupsended in 500 μl of 50 mM NaOH and one μl of the supernatant was used as a template in the following PCR reaction; plasmid libraries were diluted 10-fold in 50 mM NaOH, which was done to mimic the lysis condition used for *P. aeruginosa*, and one μl of these dilutions was used as a template in the following PCR. A 50-μl PCR reaction in GC buffer contained, in addition to the template, 0.5 μl Phusion polymerase, 200 nM dNTPs, 3% DMSO and 500 nM of forward and reverse primers each. We used a set of four “staggered” primer sets to increase sequence complexity: 16587/16591 (par0), 16588/16592 (par1), 16589/16593 (par2), and 16590/16594 (par3). PCR conditions were the following: 3 min at 98°C; 10 sec at 98°C, 10 sec at 63°C, 10 sec at 72°C (20 cycles); 1 min at 72°C; hold at 10°C. PCR products of approximately 150 bps were run on a 2% agarose gel and purified using a NucleoSpin Gel & PCR Clean-up Mini kit (MN) with elution in 40 μl ddH_2_O. PCR products were quality-checked on a AATI Fragment Analyzer (Genomics Facility Basel, ETHZ, University of Basel) and concentrations were in the range of 1-13 ng/μl. If applicable, PCR products were diluted to 1 ng/μl and the diluted DNA was used as a template in an Indexing PCR (Nextrea DNA UD Index Set C; REF: 20026934, LOT: 20580288). The Indexing PCR mix (25 μl total volume) contained 5 μl template, 5 μl primer mix, 5 μl Phusion GC buffer, 0.5 μl dNTPs and 0.25 μl Phusion polymerase. Cycling conditions were: 3 min at 98°C; 10 sec at 98°C, 10 sec at 55°C, 10 sec at 72°C (8 cycles); 1 min at 72°C; hold at 10°C. PCR products were purified using AMPure XP Beads (Beckman Coulter). Samples were submitted for Illumina sequencing to the Genomics Facility Basel (ETHZ, University of Basel) using a NextSeq SR 81/10 kit. Each sample produced around 10 million reads.

### Data Analysis of CRISPRi screens

Data analysis was performed using the European Galaxy server (Galaxy Community, 2024). First, all fastq.gz files from a single biological replicate were concatenated using the “Concatenate multiple datasets, tail-to-head by specifying how” tool, and concatenated files were saved in fastq.gz format. After an initial FASTQC check to ensure good quality of sequences, the concatenated files were analyzed using the MAGeCK pipeline (Li *et al*, 2014). sgRNAs in each sample were first counted using “MAGeCK count” using a file with all designed 83’872 as the sgRNA library reference file and the “Median” normalization method. Next, pairwise comparisons between conditions were performed using “MAGeCK test” using the manually curated “MAGeCK count” output file containing counts of all samples as input file. For all comparisons, all four biological replicates of one conditions were set as “control” or “treated” as appropriate. The following parameters were used: Method for normalization: Median; P-value adjustment method: FDR; Sorting criteria: “Negative selection”; Remove zero: Both; Remove zero threshold: 10; Gene Log-Fold Change method: “Median” or “Second-best”. From Gene Summary files, for each gene, P-values for negative and positive selection columns were compared and the smaller value was printed together with its corresponding FDR and LFC values in a new list containing only “Gene”, “P-val”, “FDR” and “LFC” columns using the AWK program. Data from this combined Gene Summary and original sgRNA Summary files were further analyzed and plotted using custom Python code. For global enrichment/depletion analyses, we only included CDSs represented by at least 3 sgRNAs with at least 10 reads per sgRNA in at least one of the two conditions. Genes with a P-value of <=0.05 and a LFC of >=1 or <=-1 were considered significantly enriched or depleted, respectively, and are included in Venn diagrams shown in Fig 2. Enriched pathways, functions or clusters in particular sets were retrieved from the String database (Szklarczyk *et al*, 2023). In the clustermap file (Table S3) all genes that are significantly enriched or depleted in at least one condition are listed along with their LFCs in all conditions tested; note that an absolute LFC of >=1 does not necessarily mean that this change was also significant in this specific condition. A sgRNA counts summary file is given as Table S2, a summary of the LFCs of all sgRNAs for all pairwise comparisons is given as Table S4, and raw data were deposited on Zenodo (https://zenodo.org/): FASTQ files 24h, 10.5281/zenodo.15591742; FASTQ files 48h, 10.5281/zenodo.15592238; MAGeCK output files (and additional data summary files), 10.5281/zenodo.15593361.

## Supplemental material

Supplementary Data 1 and Tables S1-S4 are available from Zenodo: 10.5281/zenodo.15593361. Table S5 is provided with this manuscript.

## ACKNOWLEDGEMENTS

We thank Fabienne Hamburger for help with cloning and library construction, Michael Laub, Julia Vorholt, Kristala Prather, Herbert P Schweizer and Marek Basler for plasmids; and Anne Francez-Charlot for providing Python scripts for CRISPRi analysis. This work was supported by Swiss National Science Foundation (SNSF) grant 310030_208107 and NCCR AntiResist funded by SNSF grant 51NF40_180541 to U.J. and DFG grant no. KL 3317/1 to A. Klotz.

## Notes

https://doi.org/10.5281/zenodo.15591742

https://doi.org/10.5281/zenodo.15592238

https://doi.org/10.5281/zenodo.15593361

### Competing Interest Statement

The authors have declared no competing interest.

